# An autonomous mathematical model for the mammalian cell cycle

**DOI:** 10.1101/2022.05.04.490568

**Authors:** Katherine S. Williams, Timothy W. Secomb, Ardith W. El-Kareh

## Abstract

A mathematical model for the mammalian cell cycle is developed as a system of 13 coupled nonlinear ordinary differential equations. The variables and interactions included in the model are based on detailed consideration of available experimental data. Key features are that the model is autonomous, except for dependence on external growth factors; variables are continuous in time, without instantaneous resets at phase boundaries; cell cycle controllers and completion of tasks associated with cell cycle progression are represented; mechanisms to prevent rereplication are included; and cycle progression is independent of cell size. Eight variables represent cell cycle controllers: Cyclin D1 in complex with Cdk4/6, APC^Cdh1^, SCF^βTrcp^, Cdc25A, MPF, NUMA, securin-separase complex, and separase. Five variables represent task completion, with four for the status of origins and one for kinetochore attachment. The model predicts distinct behaviors consistent with each main phase of the cell cycle. The response to growth factors shows restriction-point behavior. These results imply that the main features of the mammalian cell cycle can be accounted for in a quantitative mechanistic way based on known interactions among cycle control factors and their coupling to tasks involved in replication of DNA. The model is robust to parameter changes, in that cycling is maintained over at least a five-fold range of each parameter when varied individually. The most sensitive parameters are those associated with the initiation and completion of mitosis. The model is suitable for exploring how extracellular factors affect cell cycle progression, including responses to metabolic conditions and to anti-cancer therapies.

**Author Summary:** We created a model, that is, a set of mathematical equations, to represent the entire cell cycle in mammals. All terms in our equations correspond to actual biological mechanisms. We solved the equations and verified that they show similar behavior to real-life cell cycles. We designed our model to cycle only when enough external growth factors stimulate it, as real cells do. Our model helps us better understand how the cell cycle is controlled. One very important aspect of control is how the cell ensures that its DNA is copied only once per cell cycle. If “rereplication” (copying a section of DNA twice in the same cycle) occurs, it can cause harmful DNA damage. We considered several mechanisms that some biologists believe play a role, and found that most could not explain how rare rereplication is. Two mechanisms did work well, and we used them to make two variants of the model, which show similar behavior. The model has many potential applications. The cell cycle is altered in most cancer cells, so understanding how these changes affect control is useful. Many widely-used drugs affect the cell cycle, and our model can be used to study these effects.

## Introduction

Biological development and function depend on cell division, which is governed by the cell cycle. In human cancers, cell cycle regulation is dysfunctional, and many anti-cancer treatments disrupt the cycle. Schedule dependence of the response of cancer cells to some drugs or drug-radiation combinations has been attributed to cell-cycle disturbances caused by the therapies, as well as varying sensitivity over the cell cycle to therapy-induced damage. Improved understanding of how the cell cycle is altered in cancerous cells and how this affects cell cycle progression may lead to new therapeutic approaches. Mathematical models for the cell cycle have the potential to lead to improved quantitative understanding of cell cycle regulation, and to be useful for predicting or optimizing responses to treatments that affect the cell cycle. A model for untransformed cells can provide a basis for modeling the cell cycle in cancerous cells.

### Review of previous theoretical models

Many mathematical models for the cell cycle have been developed. Some focus on a specific phase of the cycle, such as the restriction point [1–3], the G1-S transition [3–6], DNA replication initiation [7], M phase [8], and the spindle checkpoint [9]. Some models are designed for specific cell types or experimental systems, such as the bacterium Caulobacter crescentus [10], the Xenopus frog [8,11,12], Drosophila embryos [13], yeast [7,11,14–17] and the GS-NS0 cell line, derived from mouse plasmacytoma [18]. Some models are for transformed cells, such as mouse hybridoma cells [19], HeLa cancer cells [11,20] or osteosarcoma and neuroblastoma cells [21]; cell cycle control in such cells may differ significantly from that in normal cells. Other models have been developed for pathological conditions, such as DNA damage [22,23].

In the well-known model of Tyson and Novak [24], a prescribed increase in cell size over time governed by the logistic equation drives the cell cycle and acts as the trigger for mitosis. In other models [25–28], the cycle is driven by exponential growth of cell mass. Such non-autonomous models cycle continuously with fixed cycle length, independent of extracellular signals. In reality, cells cycle only when stimulated by growth factors and when adequate nutrients are available. The non-autonomous models implicitly assume that the cell has a mechanism to detect its size and transmit this information to cell cycle controllers. There is evidence in yeast cells for a “cell-size” checkpoint [29], but the existence of such a checkpoint in mammalian cells remains controversial [30,31]. While a G1 sizer was found in mouse epidermal cells [32], it is not yet known whether this finding extends to most mammalian cells, and this sizer may not be the primary determinant of S-phase entry under adequate nutrient conditions [31]. Observations that proliferation of primary mammalian cells is independent of cell size [33], that cell growth and division are loosely coupled [34], and that growth and cell cycle progression can be influenced independently by extracellular conditions [35], suggest that cell size is not a primary trigger for mitosis or G1 exit. Regardless of cell size, cycle progression should not occur if essential tasks such as origin licensing and DNA replication are incomplete.

Fauré et al. [36] presented a Boolean model for the mammalian cell cycle, in which a constant level of Cyclin D1 causes periodic cycling. Boolean models are not well suited to investigating effects of changing extracellular conditions or simulating the action of a restriction point. Another model for mammalian cells [37] includes several hundred differential equations describing 33 proteins involved in control of the cell cycle. The complexity of this model makes it difficult to discern its key mechanisms and capabilities.

The cell cycle requires completion of several tasks, such as origin licensing and initiation, DNA replication, nuclear envelope breakdown, and kinetochore attachment. Prevention of rereplication of DNA is critical. Many of the models mentioned above describe the dynamics of a sequence of cell cycle controllers, but few models represent task completion quantitatively. The model of Chen et al. [25] includes kinetochore attachment and licensing of origins, connecting their progression to cell cycle controllers. The completion of tasks is ensured by imposing instantaneous changes in system state once a threshold is reached, and the representation of mechanisms that ensure separation between tasks is therefore simplistic. For example, instead of an abrupt shift from licensing to origin firing at the G1-S boundary, licensing is actually completed a finite time before S phase begins [38].

While models for parts of the cell cycle are useful [1–9], they cannot be combined simply as modules to represent the full cycle, because some key cell cycle controllers, such as APC^Cdh1^, are involved in more than one phase, and proper coupling between phases is critical.

### Objectives of present model

In view of the points discussed above, the objective of the present study is to develop a mathematical model for the cell cycle with the following characteristics:

- *Autonomous*. The cycle should not be driven by an external “clock.” However, dependence on external growth factors is included.
- *Continuous*. Model variables should vary continuously over the entire cycle, driven by known or plausible biological mechanisms.
- *Cell cycle phases*. The model should exhibit distinct behaviors consistent with each of the four main phases of the cell cycle (G1, S, G2 and M).
- *Dependent on task completion*. The cycle should not progress if essential tasks such as origin licensing or DNA replication are incomplete.
- *Avoids DNA rereplication*. The model should ensure that DNA rereplication is virtually non-existent.
- *Growth factor dependence*. Growth factor concentration below some level should inhibit cycling.
- *Restriction point in G1*. Beyond some point in G1, the current cycle should be independent of growth factor concentration.
- *No size checkpoint*. Cycle progression should be independent of cell size.
- *Moderate complexity*. The number of variables and equations should be minimized consistent with the above objectives, so that mechanistic insights can be obtained readily.

The model presented here represents key biological processes of the cell cycle in sufficient detail to allow prediction of responses to extracellular factors that act on specific parts of the cycle, such as metabolic conditions and anti-cancer therapies. While the model uses phenomenological representations of biological processes, it provides a coherent modeling framework for the entire cycle in which more mechanistic descriptions of individual processes could be incorporated.

## Methods

### Overview of the model

The model is formulated as a set of 13 coupled nonlinear ordinary differential equations. It is autonomous, i.e., no time-dependent driving terms are included except in the case of changing external growth factor levels. Of the 13 dynamic variables, eight represent concentrations of substances that act as cycle controllers and five represent the extent of completion of tasks, including the licensing and firing of origins, DNA replication and attachment of kinetochores. The model variables and their interactions are represented schematically in Figure 1, and are described in detail below.

**Figure 1.**
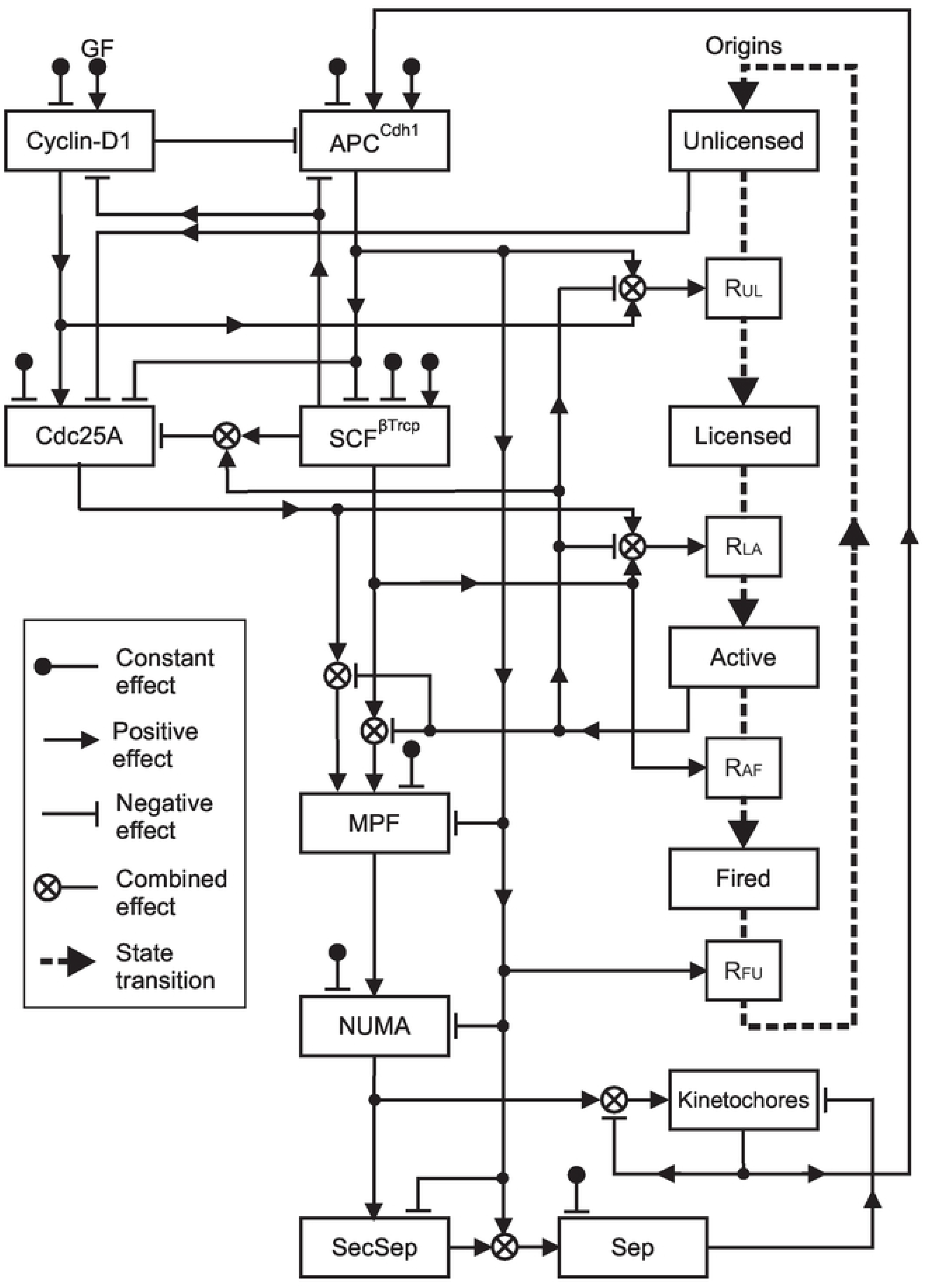
Schematic diagram of model. Model Variant 1 is shown. Rectangles denote the 13 system variables. “Cyclin-D1” is the Cyclin D1-Cdk4/6 complex. Squares and dashed lines denote transitions between states of kinetochores. Solid lines denote effects of system variables on other variables or transition rates, which may be stimulatory inputs (arrowhead) or inhibitory inputs (flat line end). Each input corresponds to a specific term in the corresponding governing equations. In the case of inhibitory inputs, the corresponding term is proportional to the current value of the affected variable.

Rapid changes in behavior occur during the cell cycle, and tasks must be completed to high levels of precision for successful cycling. Model behavior must therefore depend sensitively on system variables at some points in the cycle. To achieve this, Hill-type dependence of rates on concentrations is assumed for several processes. The exponent *n* in the Hill equation governs the steepness of the response, with *n* = 1 giving Michaelis-Menten kinetics and *n* → ∞ giving a step-function response. Available experimental data do not allow precise estimation of *n*. Here, a reference value *n* = 5 is assumed for all Hill-type responses. This value is large enough to provide the necessary sensitivity. During model development, Hill-type kinetics were assumed for multiple processes and the sensitivity of the model to each Hill exponent was tested. Michaelis-Menten or linear kinetics were then substituted if a Hill exponent greater than one was not required for satisfactory model behavior. The rationale for retaining Hill-type behaviors, where needed, is stated below.

### Choice of model variables

Many entities are involved in the cell cycle. To reduce model complexity, some sequences of steps involving several intermediaries are represented by single variables. Phosphorylated Cdc25A is taken as the main driver of S-phase entry, as it is required for S-phase entry in human cells [39–42] and is upstream of other species considered to play that role. Cdc25A phosphorylates and thus activates Cdk1 (also known as Cdc2) and Cdk2 [42], and is upstream of cyclin A/Cdk2 and cyclin E/Cdk2 [43–45], all of which have been considered as instigators of S-phase entry. Additional reasons for choosing Cdc25A over either Cdk1 or Cdk2 are that Cdk2 is not essential for cycling [46–48] and that Cdk1 and Cdk2 are mutually redundant [49,50]. Cdc25A is inhibited by deficiencies in origin licensing [51], and its use as a controller ensures that S-phase entry does not occur until adequate licensing has occurred. Cdk2 activation, downstream of Cdc25A, has also been shown to depend on origin licensing [52–54].

The inclusion of the E3 ubiquitin ligases APC^Cdh1^ and SCF^βTrcp^ as model variables instead of specific cyclins (cyclin A, cyclin E, etc.) is based on studies [55–59] identifying them as two major cell cycle controllers that alternate in dominance during the cycle through negative feedback. The anaphase-promoting complex (APC) is a cycle controller that, when bound to a coactivator protein, degrades other regulatory proteins in G1 and in late mitosis. In G1, Cdh1 is the coactivator, while in mitosis, both Cdc20 and Cdh1 associate with APC as coactivators [60]. APC in some form drives anaphase entry, which must be promoted by satisfaction of the spindle checkpoint. While APC^Cdc20^ has been considered to be the controller that responds to spindle checkpoint completion [61], other studies [62,63] point to APC^Cdh1^ as fulfilling this function, in addition to its role in G1. The level of the complex APC^Cdh1^ is therefore included as a system variable, rather than APC^Cdc20^.

The eight cell cycle controllers included in the model are the cyclin D1-Cdk4/6 complex, which promotes passage through G1; APC^Cdh1^ and SCF^βTrcp^, as discussed above; cell division cycle 25 A (Cdc25A), which is needed for the G1-S transition; mitosis promoting factor (MPF, also known as mitotic cyclin B/Cdc2 kinase, cyclin B-Cdk1 complex, or maturation promotion factor), a key driver of mitosis [64,65] responsible for breakdown of the nuclear membrane [65,66]; nuclear mitotic apparatus protein (NUMA), which promotes attachment of kinetochores to the mitotic spindle complex [67]; the securin-separase complex, which when split releases the activated enzyme separase, causing separation of sister chromatids. The model variables [CyclinD1], [APC^Cdh1^], [SCF^βTrcp^], [Cdc25A], [MPF], [NUMA], [SecSep] and [Sep] represent the concentrations of these substances in the chromosomal region, except for [NUMA], which refers to concentration at the mitotic spindle complex formed in the cytoplasm before nuclear envelope breakdown.

The five variables that quantify completion of cell cycle tasks range from 0 to 1. Kinetochore attachment (*K*) is normalized relative to 92, the total number of kinetochores. Equations are presented for two versions of the model, Variant 1 and Variant 2, depending on the assumed mechanism for prevention of rereplication, as discussed below. The status of origins is modeled by representing the number in each of several states as a fraction of the total number that are licensed during the cell cycle. The number of origins in human DNA is estimated as 50,000 to 250,000 [68], with 20,000 to 50,000 initiated in a single cycle [68,69]. The origin states are: unlicensed (*U*), partially licensed (*L*_0_, Model Variant 2 only), fully licensed (*L*), actively replicating (*A*) and fired (*F*, Model Variant 1 only), such that *U* + *L* + *A* + *F* = 1 in Model Variant 1 and *U* + *L*_0_ + *L* + *A* = 1 in Model Variant 2. “Actively replicating” refers to origins that have given rise to a replication fork that remains active. This variable (*A*) is used as a surrogate for replication protein A (RPA) which, by sensing single-stranded DNA associated with actively replicating forks, detects incomplete replication and activates the ATR-CHK1 pathway. The total amount of replicated DNA is approximately proportional to the number of fired origins. Nuclear envelope breakdown is modeled by a rapid increase in NUMA concentration at the spindle complex and formation of the securin-separase complex at the chromosomes. Cytokinesis is identified by free activated separase reaching a threshold concentration sufficient to break the attachments between sister chromatids.

### Mechanisms preventing rereplication

Avoidance of DNA rereplication is a critical property of the cell cycle [70]. Low levels of rereplication are difficult to detect experimentally [71] and may occur in normal cells [72,73], but reinitiation of even a single origin is potentially damaging [74,75]. Several potential mechanisms preventing rereplication have been proposed, as summarized in Table 1, with names assigned where needed for clarity. These mechanisms can be classified as global (acting on all origins simultaneously, Table 1A), or local (acting on origins individually, Table 1B). In this section, we discuss supporting evidence and limitations of these mechanisms, and select two as a basis for theoretical models.

**Table 1.** Proposed mechanisms for preventing rereplication.

| Organism | Mechanism |
| --- | --- |
| **A. Global mechanisms (acting on all origins simultaneously)** |  |
| *Licensing factor inactivation* |  |
| Mammalian | **Inactivation of Cdt1.** The increase in geminin around the G1-S transition inactivates the licensing factor Cdt1 [76]. Multiple mechanisms repress Cdt1 after S phase starts [77]. |
| Yeast | **Inactivation of multiple licensing factors.** High Cdk from late G1 to M inhibits several components of licensing [78-80]. |
| *Temporal separation* |  |
| Yeast | **Cdk two-threshold.** As Cdk rises, $T_L$ and $T_F$ are thresholds for inhibiting licensing and promoting origin firing, with $T_L < T_F$ [81]. |
| Mammalian | **Cyclin A two-threshold.** Analogous to the Cdk two-threshold mechanism, with Cdk replaced by Cyclin A [82]. |
| Mammalian | **Late G1 licensing inhibitor.** An unidentified inhibitor of licensing in late G1 ensures that licensing stops some time before S phase starts [83]. |
| Xenopus egg | **Cytoplasmic licensing factor (Model Variant 2).** A licensing factor present in the cytoplasm has access to nuclear DNA only in the brief period between nuclear envelope breakdown and the formation of new nuclear membranes in late mitosis. |
| **B. Local mechanisms (acting on individual origins at different times)** |  |
| *Licensing-dependent origin inactivation* |  |
| Xenopus sperm;<br>Xenopus egg | The process of licensing lowers the affinity of essential licensing factor Cdc6 for chromatin [84] and destabilizes the association of ORC and Cdc6 with chromatin, even prior to Cdk exposure [85]. |
| *Replication-dependent origin inactivation* |  |
| Fission yeast | **MPF-bound post-replicative origin state (Model Variant 1).** Origins in the post-replicative state are incapable of licensing until they are “reset” in mitosis. Specifically, MPF attaches to origins after replication and is destroyed in late mitosis by APC<sup>Cdh1</sup>. |
| Human transformed;<br>Chinese hamster | **Chromatin compaction post-replicative origin state.** Sections of replicated DNA become part of compacted/condensed chromatin [86,87] that cannot be licensed until decondensation after mitosis [88]. |
| Xenopus egg | **Replication-dependent licensing factor inactivation.** Factors such as RPA that are essential for origin initiation also destroy Cdt1, the licensing factor [89]. |
| Escherichia coli | **Origin sequestration.** After initiation, an origin is “sequestered” and unable to initiate again for approximately one-third of the normal cycle duration [90]. |
| Escherichia coli | **Regulatory inactivation of DnaA.** The association of DnaA (needed for initiation) with origin DNA is inhibited [91]. |
| Escherichia coli; fission yeast | **Titration of DnaA.** As DNA is replicated, the new DNA avidly binds DnaA around the origin site, making it unavailable for further initiation, as it is not produced during S phase [92]. |

According to *Licensing factor inactivation*, prevention of rereplication depends on inactivation of factors required for licensing of origins, such as the replication licensing factor Cdt1. The nuclear protein geminin inactivates Cdt1 [76], and depletion of geminin with consequent overexpression of Cdt1 was found in transformed cells to lead to rereplication [93]. However, overexpression of Cdt1 in the untransformed human fibroblast cell line HFF-2 does not lead to rereplication [94]. Also arguing against this mechanism is that measurable Cdt1 levels are observed in S phase [95,96].

An alternative theory, *Inactivation of multiple licensing factors*, is that a single factor inhibits multiple licensing factors [78,79,97]. In yeast, for example, Cdk deactivates or degrades several components required for licensing. If the probability of licensing depends on the product of the concentrations of several factors, which are all reduced to very low levels, then the resulting probability would be virtually zero [97]. However, Cdk activity is reduced through inhibition of Cdc25A in the DNA damage response [80], and would therefore be ineffective in preventing rereplication in the case of DNA damage. This suggests that Cdk-independent mechanisms for prevention of rereplication prevention are likely to be more important.

Mechanisms of this type have a theoretical weakness [75]. If a transition between phases is governed by an exchange between two mutually inhibiting factors, the second factor (e.g. Cdt1) must rise to a sufficient concentration to cause inhibition, before the concentration of the first one (e.g. geminin or Cdk1) drops to a low level, causing a period of overlap when both concentrations are nonzero. The same limitation applies to the *Licensing-dependent origin inactivation* mechanism (discussed below).

Several mechanisms based on temporal separation have been proposed as ensuring that origin licensing ends before origin initiation begins [81–83]. In two-threshold models, a single factor (Cdk or Cyclin A) inhibits origin licensing when one threshold is reached and permits origin firing when a second higher threshold is reached. Such a mechanism relies on consistency of thresholds across cell populations, as well as sensitive dependence of rates of licensing and firing on concentration, in order to achieve complete temporal separation, and therefore lacks robustness.

According to the *Cytoplasmic licensing factor* mechanism, an essential factor for licensing is present in the cytoplasm, and reaches DNA only in the period between nuclear envelope breakdown and reformation of the nuclear envelope in mitosis, allowing the first step of origin licensing to occur in mid/late M, with the licensing process completing in G1. In Xenopus eggs, evidence was found for a licensing factor that cannot cross the nuclear membrane to enter the nucleus [98,99]. In transformed human cell lines, nearly all of the key origin license component Cdc6 was found to be bound to chromatin within minutes of nuclear breakdown [100]. Observations of changes of localization during the cell cycle of Cdc6 in yeast [101] combined with findings that it cannot cross the nuclear membrane in Xenopus eggs [102] are consistent with this having a role in preventing rereplication. This mechanism is included here as “Model Variant 2.”

Local mechanisms also contribute to prevention of origin re-initiation [103]. For example, in *Licensing-dependent origin inactivation* [84,85] the final steps of licensing decrease the ability of ORC and Cdc6 (two essential components of the pre-replication complex) to remain attached to chromatin.

The term *Replication-dependent origin inactivation* [104] was used for a mechanism in which the process of replication destroys the licensing factor Cdt1 [89]. Here it is applied more generally to mechanisms in which an altered state of origins after replication makes them resistant to re-licensing or re-initiation until a later phase in the cell cycle.

One such mechanism, suggested by a combination of studies, is termed *MPF-bound post-replicative origin state*. In a study of yeast, Dahmann et al. [105] proposed that “S-phase-promoting cyclin B–Cdk complexes prevent rereplication during S, G2 and M phases by inhibiting the transition of replication origins to a pre-replicative state.” They note that G1 but not G2 nuclei can be induced to prematurely enter S phase and suggest that differences in the pre- and post-replicative state of origins are responsible [105]. Wuarin et al. [106] found that in fission yeast, MPF associates with the post-replicative complex and prevents re-formation of the pre-replicative complex in the current cycle, and proposed that this explains why G1 nuclei can be stimulated to replicate but G2 cannot. They found that the rise of APC^Cdh1^ in anaphase removes MPF from the post-replicative complex, restoring it to the unlicensed state. This study did not directly demonstrate that MPF associates to each origin as replication is completed. However, a mutually exclusive association of MPF and minichromosome maintenance protein complex (MCM) with replication origins was demonstrated. MCM is part of the pre-replicative complex that is removed after firing, implying that that MPF could potentially associate immediately with each origin after it has fired. This mechanism is included here as “Model Variant 1.”

A second mechanism of this type is the *Chromatin compaction post-replicative origin state*, based on the following findings: chromatin compaction status determines accessibility to factors needed for DNA replication [107]; licensing and replication occur on decondensed chromatin [88,108]; newly replicated DNA immediately adopts more compacted forms or moves to areas of condensed chromatin [86,87]; compaction continues to increase in G2/M. Immediately after mitosis, decompaction of chromatin occurs [88] allowing licensing again. Mathematically, this model would be nearly identical to one using *MPF-bound post-replicative origin state*. In both cases, the states of origins are reset in late mitosis, the difference lying only in the timing and the agent triggering the reset. Four additional proposed mechanisms for replication-dependent origin inactivation, based on observations in non-mammalian systems, are listed in Table 1.

The critical necessity for cell division to occur without rereplication suggests that multiple mechanisms may be involved. The relative importance of the mechanisms is not known. Based on the preceding discussion, two mechanisms are well supported by experimental evidence and could prevent rereplication during the period where it would be detrimental (S, G2, and early mitosis), allowing resumption of origin licensing in mid to late mitosis. For these two mechanisms, *MPF-bound post-replicative origin state* and *Cytoplasmic licensing factor*, corresponding Model Variants 1 and 2 are presented.

### Derivation of model equations

In the following, the governing equation for each model variable is given and each term in the right hand side is justified based on available information about the factors influencing its dynamics.

#### Cyclin D1-Cdk4/6 complex

The equation is:

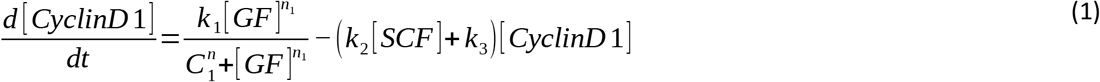

Cyclin D1 plays a key role in progression through G1 [109] and the progression from G1 to S [110], increasing in G1, declining on entering S phase [111], but remaining in G2 [112]. *Term 1:* Growth factors (mitogens), with concentration [GF] in the extracellular space cause the levels of Cyclin D1, in particular in its activated form in complexes with Cdk4/6, to increase [113–115]. The Hill equation is used to ensure that the restriction point in G1 is insensitive to growth factor concentration above a threshold value. *Term 2:* SCF^βTrcp^ degrades Cyclin D1 [111,113,116]. *Term 3:* Cell-cycle-independent degradation is assumed. The half-life of free or Cdk4-bound Cyclin D1 is less than 30 minutes [113].

#### APC^Cdh1^

The equation is:

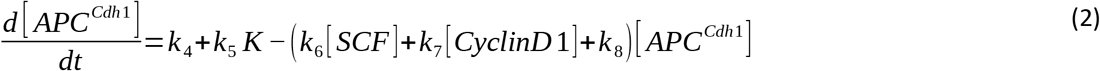

*Term 1:* Cell-cycle-independent production. *Term 2:* Satisfaction of the spindle checkpoint occurs when all 92 kinetochores are attached, so that *K* reaches 1. This releases a block on APC^Cdh1^ [62], causing an abrupt rise in its levels because kinetochore attachment is rapid. *Term 3:* SCF^βTrcp^ and APC^Cdh1^ alternate in dominance during the cell cycle [55,56,117] and degrade each other. This term represents the degradation of APC^Cdh1^ by SCF^βTrcp^ [55,60]. *Term 4:* Degradation of APC^Cdh1^ by the Cyclin D1-Cdk4/6 complex is supported by observations that Cdk4/6 induces Emi1, which in turn suppresses APC^Cdh1^ activity [118]. *Term 5:* Cell-cycle-independent degradation.

#### SCF^βTrcp^

The equation is:

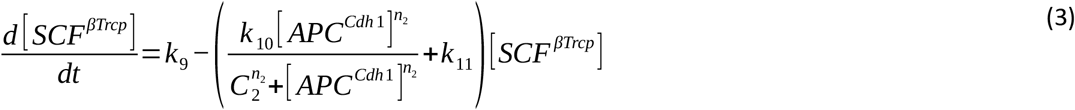

*Term 1:* Cell-cycle-independent production. *Term 2:* APC^Cdh1^ degrades SCF^βTrcp^ [55]. The Hill equation ensures that the rise in SCF^βTrcp^ occurs only when APC^Cdh1^ has reached a very low level, corresponding to the initiation of S phase. *Term 3:* Cell-cycle-independent degradation.

#### Cdc25A

The equation for Model Variant 1 is:

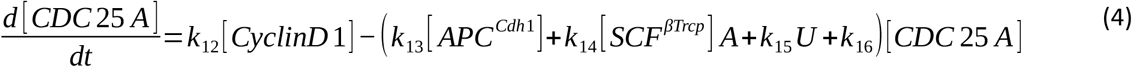

In Model Variant 2, the quantity *U* in the last term is replaced by *L*_0_. Cdc25A is important for cell cycle control [119,120], triggering S phase entry [42,121–123] when the cell has both sensed adequate growth factors over a sufficient period and achieved adequate licensing. *Term 1:* Cyclin D1-Cdk4/6 complexes promote increase of Cdc25A [124]. *Term 2:* Inhibition of Cdc25A by APC^Cdh1^ [125,126]. *Term 3:* Cdc25A is degraded by SCF^βTrcp^ when the ATR-CHK1 pathway is activated by Replication Protein A binding singlestranded DNA [42,119,127–130]. In a normal unperturbed cell cycle, single-stranded DNA is present when forks are actively replicating [131,132], i.e. when A is nonzero. While the ATR-CHK1 pathway is usually mentioned in connection with DNA damage or stalled forks, several studies have shown that it is also activated in unperturbed S phase [127,133,134]. *Term 4:* Deficiencies or defects in origin licensing cause strong inhibition of Cdc25A [51,135]. In Model Variant 1, this is represented by *U* not being close to zero, and in Model 2 by *L*_0_ not being close to zero. *Term 5:* Cell-cycle-independent degradation.

#### U, number of unlicensed origins

The equations for Model Variants 1 and 2 are:

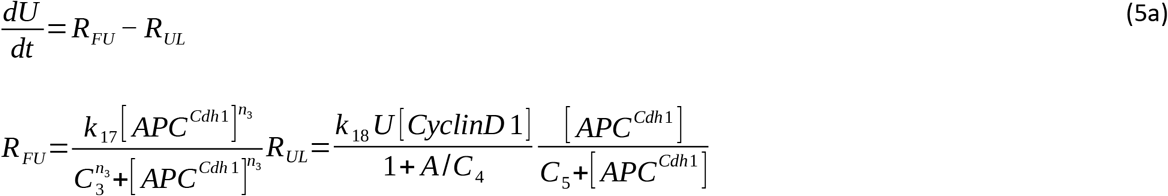

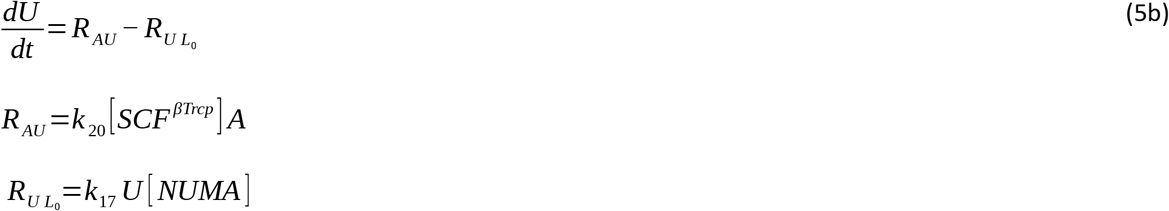

In Model Variant 1, origins progress from unlicensed (*U*) to licensed (*L*), to actively replicating (*A*), to fired (*F*), still in the post-replicative state, with completed replication forks. Model Variant 2 includes a partially licensed pool (*L*_0_) between *U* and *L*. In Model Variant 1, the *F* pool remains until it returns to the pre-replicative unlicensed state (*U*) at anaphase. In Model Variant 2, the origins with actively replicating forks (*A*) go directly into *U* when these forks terminate. *Term 1*. In Model Variant 1, *R_FU_* is the rate at which fired origins are restored to the unlicensed state, triggered by the increase of APC^Cdh1^ at the end of mitosis. A Hill equation is used here to prevent premature resetting at low levels of APC^Cdh1^. In Model Variant 2, *R_AU_* is the rate at which origins with terminated forks are restored to the unlicensed state. The dependence on SCF^βTrcp^ arises from metabolic considerations. The rate of new DNA production depends on the levels of glutamine and glutaminase [136]. Nutrient levels are assumed to be adequate in the present model, so glutamine is not modeled explicitly. SCF^βTrcp^ is used here as a surrogate measure for glutaminase, because glutaminase is inhibited by APC^Cdh1^ [58,137], which is low when SCF^βTrcp^ is high. *Term 2*. In Model Variant 1, *R_UL_* is the rate of licensing, which is damped by the number *A* of origins with actively replicating forks [89,138]. As mentioned above, the ATR-CHK1 pathway is activated by Replication Protein A binding single-stranded DNA that is present at active forks in unperturbed S phase [127,133]. This pathway controls S phase by inhibiting firing of new origins.

Cyclin D1-Cdk4/6 is required for licensing as it dissociates RB-MCM7 complexes, a necessary step in formation of the pre-replicative complex [139]. The rate of licensing has a saturating dependence on APC^Cdh1^ concentration because the licensing factor Cdt1, needed for pre-replicative complex formation, is present only when APC^Cdh1^ is at significant levels. APC^Cdh1^ prevents buildup of geminin. When APC^Cdh1^ levels go down, geminin increases and forms an inhibitory complex with Cdt1, preventing licensing [140]. In Model Variant 2, *R*_*U L*_0__ is the rate at which unlicensed origins move into the partially licensed pool (*L*_0_). This rate is assumed proportional to [NUMA], which is used here, for model parsimony, as a surrogate variable for the concentration of cytoplasmic licensing factor, possibly Cdc6, in the chromosomal region. Cdc6 is restricted to the cytoplasm when the nuclear membrane is present, but has access to origins once the membrane is dismantled. [NUMA] and [Cdc6] both rise on nuclear envelope breakdown, although [NUMA] is in a different spatial location. The evidence for this was discussed in the context of the Cytoplasmic Licensing Factor model for preventing rereplication.

#### L_0_, number of partially licensed origins

The equation (Model Variant 2 only) is:

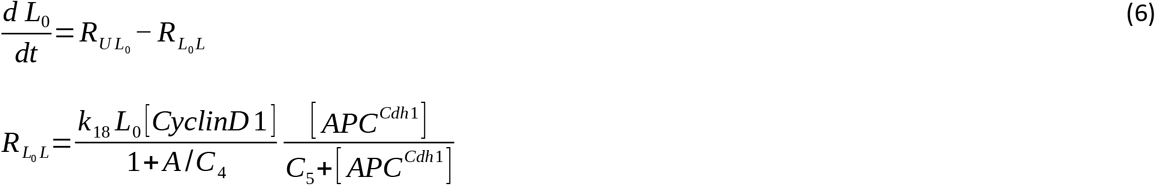

*Term 1*. *R*_*U L*_0__ was discussed above. *Term 2:* In G1, origins that were partially licensed in late M become fully licensed. This corresponds to the term *R_UL_* in Model Variant 1.

#### L, number of fully licensed origins

The equations for Model Variants 1 and 2 are:

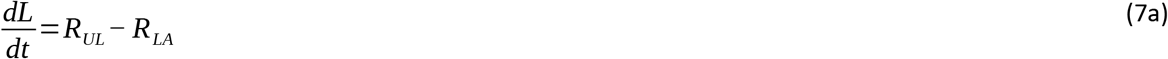

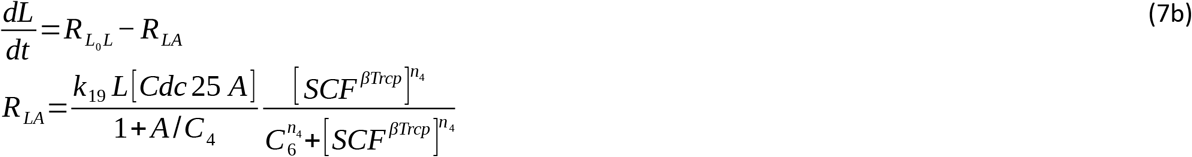

*Term 1*. The first term in both variants is the rate of G1 licensing, as already discussed. *Term 2*. In both variants, *R_LA_* represents the rate of origin firing (or initiation). This requires Cdc25A, which is needed for S phase entry [42,121–123]. A damping factor depending on the number of actively replicating origins *A* is included. Active replication forks have single-stranded DNA [131] which binds Replication Protein A, activating the ATR-CHK1 pathway and inhibiting DNA replication. In the absence of any damping, replication of the human genome would take about an hour under optimal nutrient conditions, based on a maximal fork velocity 2 kb/min [136] and an average inter-origin spacing (replicon size) of 125 kb [141], whereas S phase in a normal human cell cycle takes 8-10 hours [142]. The final factor, which depends on SCF^βTrcp^, is used as a surrogate measure for glutaminase, i.e., glutamine metabolism, as discussed above. The Hill equation is needed here to achieve separation between licensing and firing of origins.

#### A, number of origins with actively replicating forks

The equations for Model Variants 1 and 2 are:

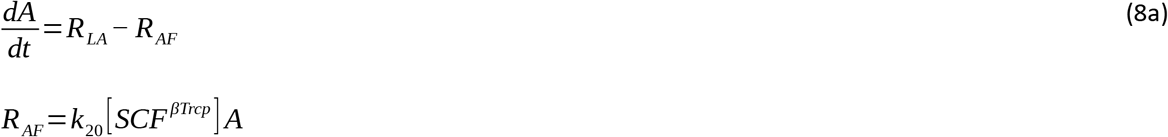

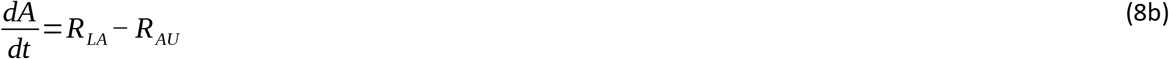

*Term 1*. The first term in both variants is the rate of origin firing, as already discussed. *Term 2* in the two model variants represents the rate at which origins cease to have active forks, and enter the fired or unlicensed pools, respectively. This rate is also a measure of the rate of DNA replication.

#### F, number of fired origins whose associated forks have terminated

The equation (Model Variant 1 only) is:

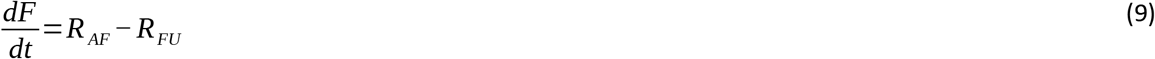

#### Mitotic-promoting factor (MPF)

The equation is:

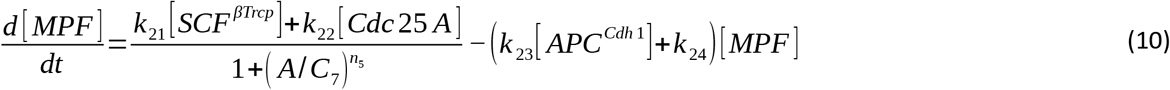

*Term 1:* SCF^βTrcp^ promotes (cyclin B-Cdk1 complex) MPF by targeting Cdk1 (CDC2) inhibitors such as Wee1 [143] and p21 and p27 [144]. However, Replication Protein A-bound ssDNA (single-stranded DNA) activates the CHK1 pathway which inhibits Cdk1 [145], so that MPF is blocked until DNA is completely replicated and there is no more RPA-bound ssDNA. Here, *A* is used as a surrogate measure of amount of ssDNA. The factor in the denominator keeps the rate low until *A* is close to zero. *Term 2:* Cdc25A promotes MPF [42,146]. T*erm 3:* APC^Cdh1^ degrades cyclin B [147,148]. MPF, which is a complex containing cyclin B, must be degraded before mitotic exit. Before APC^Cdh1^ rises again in the latter part of mitosis, APC^Cdc20^ may perform some of this degradation [148,149] but here it is attributed primarily to APC^Cdh1^. *Term 4:* Cell-cycle-independent decay is assumed.

#### NUMA (at the spindle complex)

The equation is:

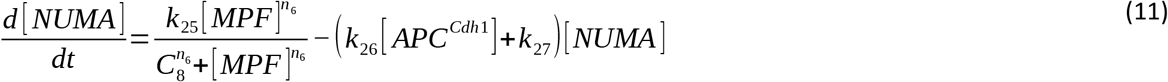

*Term 1:* [MPF] rises until it triggers nuclear envelope breakdown, which occurs on a timescale of minutes [150,151]. On nuclear envelope breakdown, NUMA from the nuclear compartment can reach the spindle complex, which was formed in the cytoplasm [152]. This is modeled as an abrupt rise in NUMA concentration at the spindle complex driven by MPF, resulting from the assumed Hill-type kinetics. *Term 2:* APC^Cdh1^ destroys mitotic proteins and cyclins towards the end of mitosis [153]. *Term 3:* Cell-cycle-independent decay is assumed.

#### K, number of attached kinetochores

The equation is:

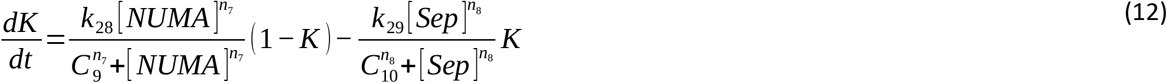

*Term 1*. After nuclear envelope breakdown, NUMA stimulates kinetochore attachment to the spindles, at a rate proportional to the number of unattached kinetochores. *Term 2*. The detachment of kinetochores at the end of mitosis is assumed to be driven by the rise in separase. In both terms, the Hill equation is used to represent abrupt rise and fall of kinetochore attachment.

#### Securin-separase complex [SecSep]

The equation is

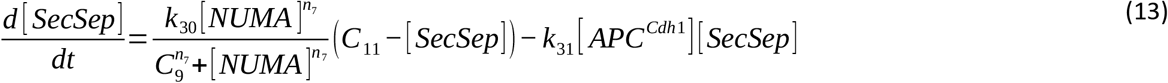

The enzyme separase cleaves cohesin, releasing the bond that holds sister chromatids together [154]. Premature targeting of cohesin by separase in early mitosis is believed to be prevented by the formation of securin-separase, an inactive complex. In late mitosis, APC^Cdh1^ cleaves this complex [62,63], releasing activated separase [154]. Jallepalli et al. [155] found that securin is necessary for chromosomal stability in human cells, and contributes to proper separase function. Other reports [156,157] imply that securin is not essential for successful cell cycling in mammalian cells, indicating that other mechanisms contribute to stabilizing separase. The form of the equation does not depend on the specific entity that stabilizes separase. Formation of separase in the cytoplasm is not represented in the model. Nuclear [SecSep] is assumed to be promoted by NUMA, which here is functioning as a surrogate variable for the amount of free unactivated separase in the chromosomal region. Separase is in the cytoplasm and excluded from the nucleus before nuclear envelope breakdown, and only once it reaches the chromosomal region can it form the [SecSep] complex with securin. NUMA is an appropriate surrogate variable for this free unactivated separase because both it and NUMA rise simultaneously due to nuclear envelope breakdown, even though they are in different spatial regions. Note that this free unactivated separase is not included in the model variable [Sep] which only represents activated separase, that is, separase released from [SecSep]. The same Hill function as in Eq. (12) is assumed. *C_11_* represents the amount available from the cytoplasm. *Term 2*. Cleavage of the complex by APC^Cdh1^.

#### Free activated separase [Sep]

The equation is:

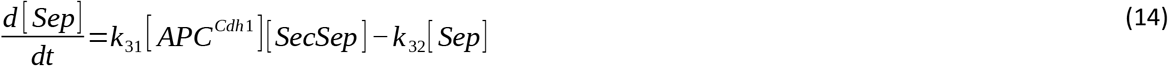

*Term 1*. Separase first forms a complex with securin and is then released in active form, targeting cohesin and promoting chromatid separation. Cleavage of the securin-separase complex by APC^Cdh1^ releases activated separase [154,155,158,159]. *Term 2:* Cell-cycle-independent decay is assumed. *Cytokinesis*. Chromatid separation is assumed to be triggered when free activated separase rises to a threshold level[*Sec*]_*th*_ sufficient to target cohesin:

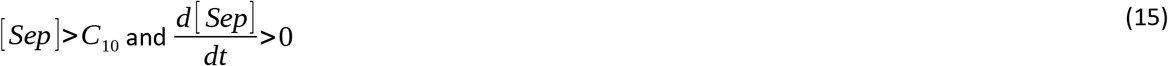

Cell cycle period is defined as the interval between these events.

The model contains 52 parameters, with reference values given in Table 2. All concentrations are considered to be dimensionless, as are the constants *C_i_* and *n_i_*. The constants *k_i_* have units h^-1^. The aim of this work is to demonstrate that the model incorporating the selected variables and mechanisms is capable of meeting the objectives stated in the Introduction. Insufficient data are available for formal parameter identification, and no claim is made that the parameter values are accurate. As discussed above, all Hill coefficients are set to 5. The reference parameter values were adjusted so that all variables (except *A*) reach a maximum of approximately 1 during the cycle, and the periods of the main cell phases agree approximately with typical values for rapidly cycling human cells, for which the overall period is about 24 hours and the G1, S, G2 and M phases last 11, 8, 4 and 1 h respectively [160]. The initial conditions are all set to zero except [APC^Cdh1^] = 1, and *U* = 1 (Model Variant 1) or *L_0_* = 1 (Model Variant 2). If periodic cycling is achieved, the eventual behavior is independent of the initial conditions. The system was solved using the ode15s routine in MatLab (MathWorks, Natick, MA). Sensitivity analysis was performed for all parameters to determine the range of values that showed cycling.

**Table 2.** Reference parameter values (Model Variants 1 and 2)

| Parameter | Value | Parameter | Value | Parameter | Value | Parameter | Value |
| --- | --- | --- | --- | --- | --- | --- | --- |
| [GF] | 1 | $k_{14}$ | 20 | $k_{28}$ | 50 | $C_1$ | 0.25 |
| $k_1$ | 1 | $k_{15}$ | 10 | $k_{29}$ | 50 | $C_2$ | 0.1 |
| $k_2$ | 2 | $k_{16}$ | 1 | $k_{30}$ | 2 | $C_3$ | 0.25 |
| $k_3$ | 1 | $k_{17}$ | 100 | $k_{31}$ | 2 | $C_4$ | 1 |
| $k_4$ | 0.02 | $k_{18}$ | 1 | $k_{32}$ | 1 | $C_5$ | 0.1 |
| $k_5$ | 1.5 | $k_{19}$ | 1.5 | $n_1$ | 5 | $C_6$ | 0.1 |
| $k_6$ | 5 | $k_{20}$ | 2 | $n_2$ | 5 | $C_7$ | 0.001 |
| $k_7$ | 0.18 | $k_{21}$ | 2 | $n_3$ | 5 | $C_8$ | 1 |
| $k_8$ | 0.025 | $k_{22}$ | 0.1 | $n_4$ | 5 | $C_9$ | 0.5 |
| $k_9$ | 1 | $k_{23}$ | 1 | $n_5$ | 5 | $C_{10}$ | 0.5 |
| $k_{10}$ | 2 | $k_{24}$ | 1 | $n_6$ | 5 | $C_{11}$ | 2 |
| $k_{11}$ | 1 | $k_{25}$ | 10 | $n_7$ | 5 | | |
| $k_{12}$ | 2 | $k_{26}$ | 2 | $n_8$ | 5 | | |
| $k_{13}$ | 10 | $k_{27}$ | 1 | | | | |

## Results

### Model representation of the cell cycle

Figure 2 shows the time course of the model variables during one cycle, for the reference parameter values and Model Variant 1. The growth factor level is held constant. Transients from the initial conditions decay after a few cycles, after which the behavior is periodic. The system of equations autonomously generates a sequence of behaviors corresponding closely to events occurring during the cell cycle.

**Figure 2.**
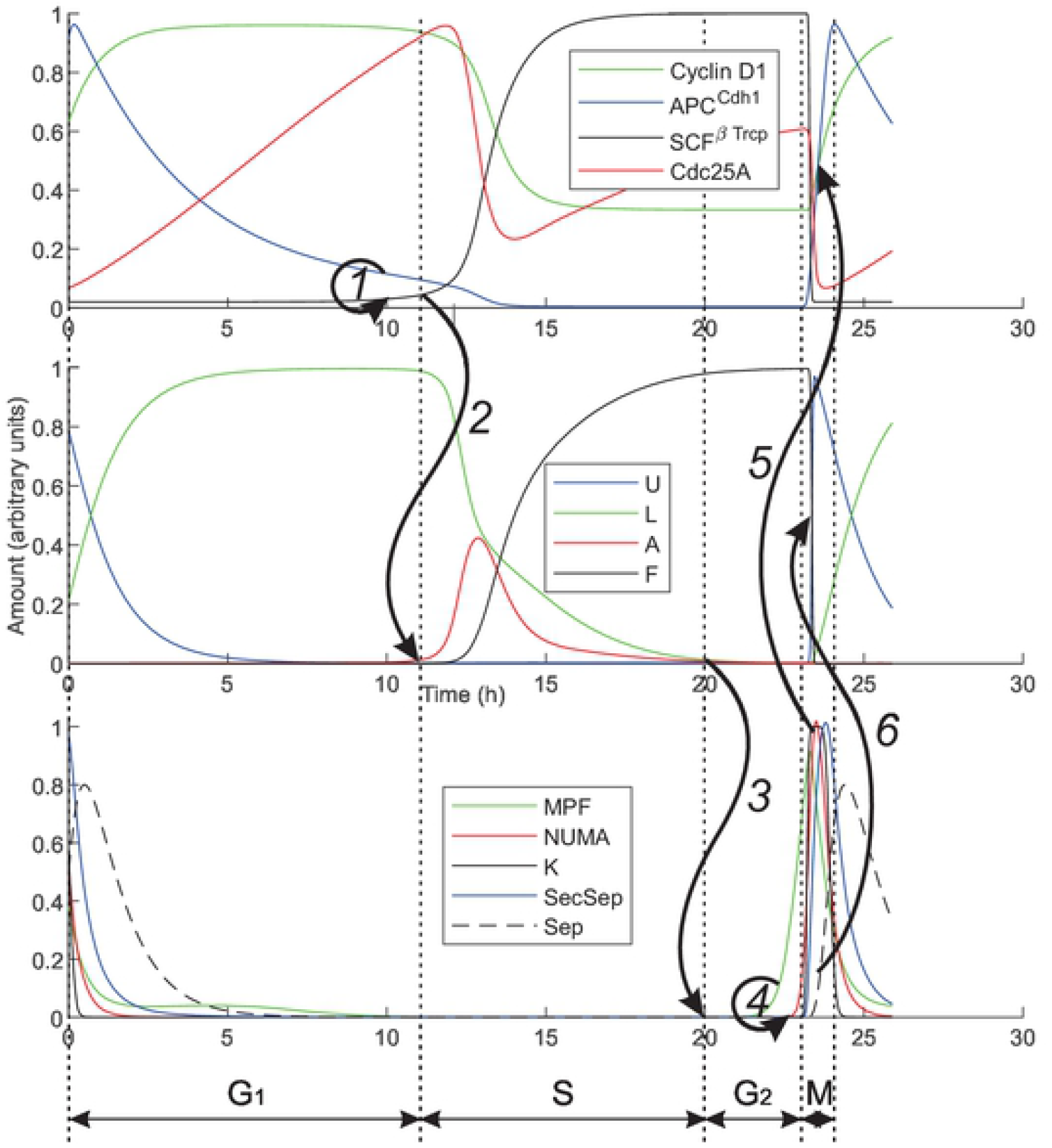
Simulated cell cycle. Time-dependent variation of system variables, showing key events involving rapid transitions in behavior. ***1***. Decreasing level of APC^Cdh1^ relieves inhibition of SCF^βTrcp^ (Eq. 3). ***2***. Increasing level of SCF^βTrcp^ triggers activation of origins (Eq. 7). ***3***. Level of *A* decreasing to a very low level, representing elimination of single-strand DNA, relieves inhibition of MPF production (Eq. 10). ***4***. Rising level of MPF triggers nuclear envelope breakdown, allowing NUMA to reach the mitotic spindle complex region and separase to reach the chromosomal region. NUMA then stimulates kinetochore attachment, and the securin-separase complex is formed at the chromosomes (Eqs. 11, 12 and 13). ***5***. Complete attachment of kinetochores triggers rapid rise in APC^Cdh1^ (Eq. 2). ***6***. Increasing APC^Cdh1^ cleaves securin-separase complex, causing a rise in separase concentration (Eq. 14). When this concentration reaches its threshold level, origins are reset to the unlicensed state and kinetochores are detached.

During the first 11 h, corresponding to G1 phase, the level of cyclin D1-Cdk4/6 rises initially and then remains high. Initially [APC^Cdh1^] is high, and the combination of high [Cyclin D1] and high [APC^Cdh1^] drives the licensing of origins. The high [Cyclin D1] causes a gradual decrease in [APC^Cdh1^] and an increase in [Cdc25A]. When [APC^Cdh1^] reaches a low level, the inhibition of SCF^βTrcp^ is released and [SCF^βTrcp^] increases. This event represents the G1-S transition and triggers the initiation of the origins.

During the next 9 h, corresponding to S phase, [SCF^βTrcp^] continues to increase, driving the firing of all origins, which remain active for some time before transiting to the fired and completed state. In this model, the fraction *F* of fired and completed origins is a surrogate measure of the fraction of DNA replicated. The sigmoidal variation of this variable during S phase is consistent with experimental data [161,162]. The production of MPF is strongly inhibited by the presence of origins with actively replicating forks. When the variable *A* reaches a value very close to zero (*C*_7_ = 0.001), the S-G2 transition is reached and MPF production can begin. The G2 phase lasts 3 h in this simulation. Toward the end of this phase, [MPF] reaches a critical level that triggers a rapid increase in [NUMA]. This represents the G2-M transition.

The constants of the model are chosen to simulate a relatively rapid sequence of events occurring during M-phase, which lasts 1 h. The rapid increase in [NUMA] upon nuclear envelope breakdown stimulates kinetochore attachment, and the variable K quickly approaches 1, representing complete attachment and satisfying the spindle checkpoint. This stimulates a rapid increase in [APC^Cdh1^]. At the same time, the breakdown of the nuclear membrane allows separase to enter the chromosomal region, where the increasing level of APC^Cdh1^ causes release of activated separase from the securing-separase complex. The rise of free activated separase triggers the events of cytokinesis, including the detachment of the kinetochores and resetting of the fired origins to the unlicensed state, completing the cycle.

### Restriction point behavior

Figure 3 shows the effects of varying the level and period of exposure of extracellular growth factor (GF) during G1 on the simulated progression through one complete cell cycle. The minimum level for cycling in the case of continuous exposure is [GF] = 0.24. The minimum level increases for shorter exposure times, but the cycle is not completed if the period of exposure is less than 9.6 hours, independent of [GF]. The results are not consistent with the proposal [163] that passage of the restriction point is dependent simply on the time-integral of growth regulatory signals.

**Figure 3.**
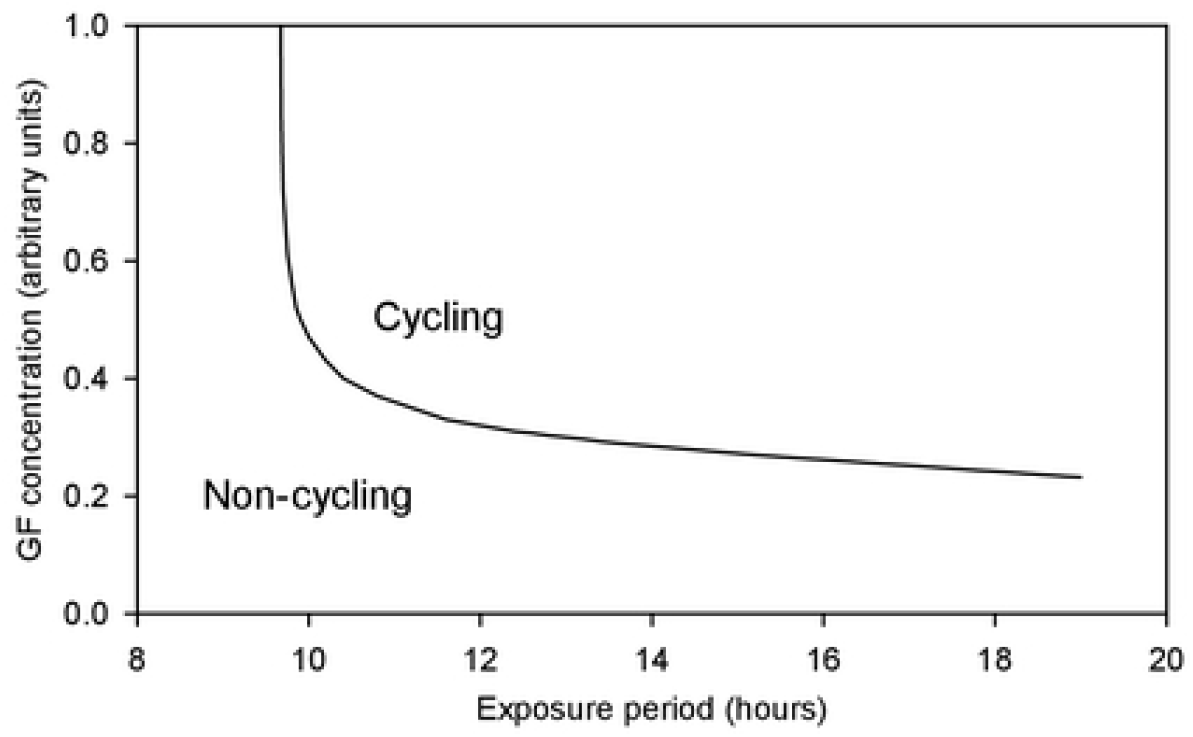
Restriction point behavior. The conditions allowing completion of one cell cycle are shown in terms of the values of *G*, representing the level of external growth factors and *t_G_*, representing the period of exposure, from the start of G1.

### Parameter sensitivity

Results of the sensitivity analysis are summarized in Figure 4. The values of the parameters were varied one at a time in small steps from 0.2 to 5 times their reference values. Vertical bars indicate the ranges of values for which a cycle was completed within 100 hours. All parameters could be varied either up or down by a factor of 5 without causing cessation of cycling. This result indicates that the cycling behavior of the model is robust, in that it is not restricted to narrow ranges of parameter values. However, model behaviors such as the overall period, the periods of the phases, the sharpness of transitions between phases, and the degree of completion of origin firing and kinetochore attachment are affected by the assumed parameter values.

**Figure 4.**
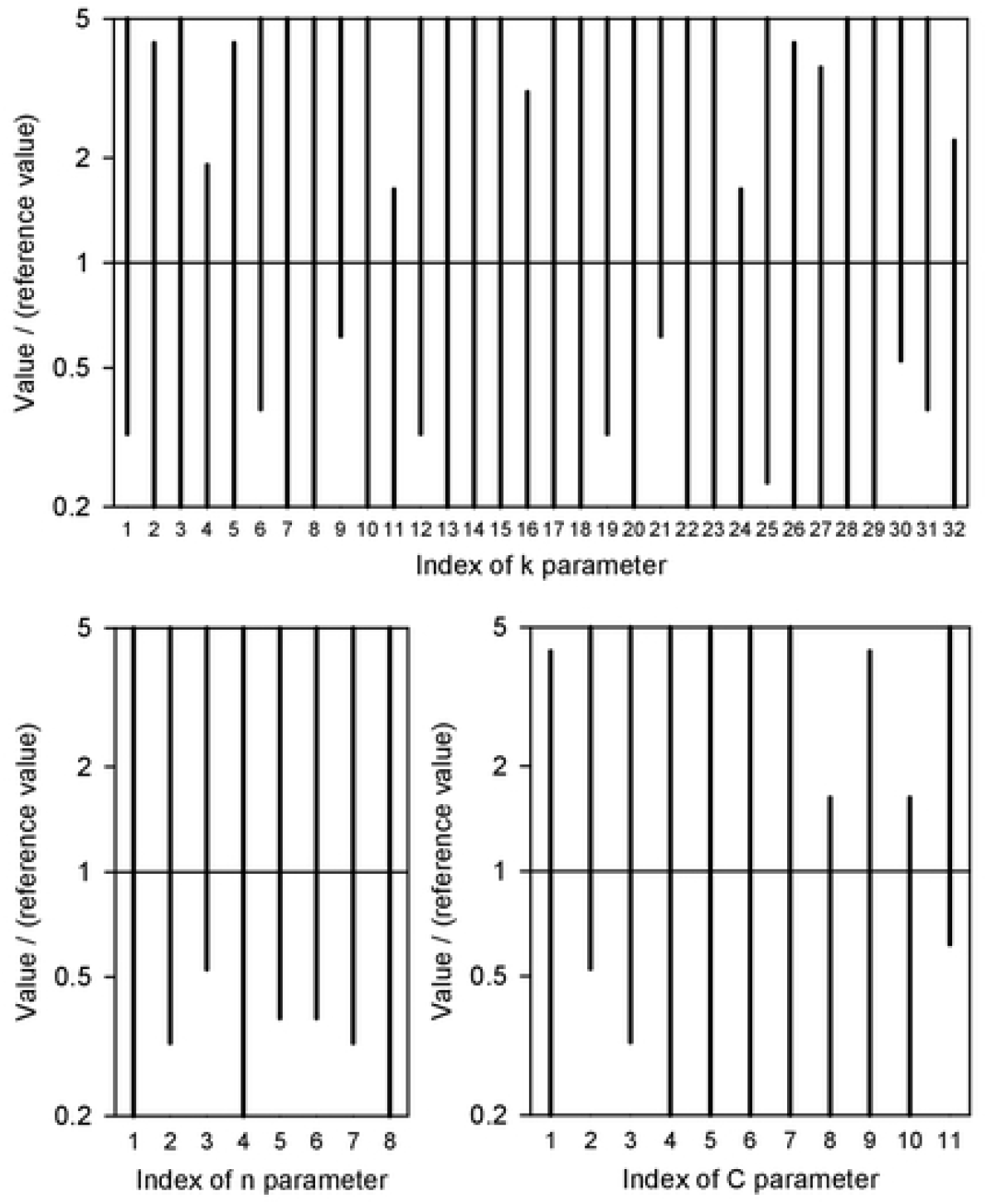
Parameter sensitivity. Results for Model Variant 1 are shown. Each parameter *k_i_* (*i* = 1,…,32), *n_i_* (*i* = 1,…,8) and *C_i_* (*i* = 1,…,11) was varied from 0.2 to 5 times its reference value, with the other values held fixed. Vertical bars indicate range of values for which cycling was completed with 100 hours.

### Model Variant 2

The schematic diagram and results for Model Variant 2, which assumes the *Cytoplasmic licensing factor* mechanism for preventing rereplication, are given in the Supplementary Material. The results are very similar to those for Model Variant 1, with the exception that the origins transit from partially licensed to fully licensed in G1, and from active to unlicensed in G2.

## Discussion

A key feature of the present model is that the equations are largely based on known biological processes and interactions. The fact that the model can reproduce the main features of the cell cycle supports the contention that these features can be understood in terms of known biological mechanisms. With 13 dynamic variables, the model has moderate complexity, and represents an approximation and simplification in many respects. Nonetheless, it provides a quantitative framework for integrating a large body of experimental knowledge about cycle mechanisms, and for investigating how these mechanisms contribute to cycle progression. For instance, the effect of a drug that interferes with the progress of a specific part of the cell cycle can be represented by including an additional factor in the differential equation representing that part of the cycle.

The process of developing this model provided a critical test of a number of proposed mechanisms for the control of the cell cycle. For example, theoretical considerations guided the selection of *MPF-bound post-replicative origin state* and *Cytoplasmic licensing factor* as potential mechanisms for prevention of rereplication. Similarly, the need to detect incomplete DNA replication led to the identification of Replication Protein A as a likely candidate, represented in the model by the number of active origins (*A*) as a surrogate measure.

The model demonstrates the possibility of a functioning cell cycle even without direct detection of DNA replication completion. The intra-S phase checkpoint mechanism, halting S-phase exit as long as any replicons remain incompletely replicated, does not preclude entry into G2 and M without any replication having taken place [164]. At present, no additional checkpoint is known that prevents progression past S phase in such a case.

Another key feature of the model is the tight integration between the activity of cell cycle controllers and the progress of essential tasks in the cycle, whose status is represented by the variables *U, L, A, F* and *K* in Model Variant 1. As indicated in Figures 1 and 2, not only are the dynamics of these variables driven by the cell cycle controllers, but the status of these variables affects the progression of the controllers. Such a two-way interaction is needed to ensure task completion, which is essential for successful cell division.

Qualitative descriptions of cell cycle control often refer to factors being present or activated in some phases and absent or inactivated in others. However, biochemical processes do not allow an instantaneous drop of concentration from a finite value to zero. In the present model, all variables show continuous variation with time. They may show rapid exponential decay during some phases, but they are never precisely zero (except at *t* = 0). This is consistent with experimental evidence. For example, the licensing factor Cdt1, often said to be absent in S phase, is present at very low levels [101,165]. The cell cycle controller DDK, stated to be “inactive” in G1 [166], actually shows some function [167]. The present model demonstrates that key functional behaviors of the cell cycle can be represented by a system in which a strict dichotomy between presence and absence of factors is not imposed.

The present model includes two types of mechanisms that permit rapid transitions in concentrations. One is the assumed Hill-type rate terms with exponents significantly higher than 1. The other is nuclear envelope breakdown and re-formation, which are rapid biomechanical processes that facilitate abrupt transitions in concentrations using a physical barrier. In the present model, nuclear envelope breakdown brings about three abrupt changes that are essential for cell cycle control: it allows cytoplasmic separase to reach the chromosomal area; it allows the cytoplasmic licensing factor to reach the chromosomal area; it allows NUMA from the nucleus to reach the spindle assembly complex region and promote kinetochore attachment.

The development of an autonomous model was motivated in part by the goal of investigating the effect of changing extracellular conditions on the progression of the cell cycle. This capability is demonstrated by the results showing the effects of varying levels and exposure lengths of growth factor (Figure 3). The model represents the G1 restriction point, in that exposure to growth factor must extend for at least 11 hours to enable progression to S phase. Beyond this point, the cycle will proceed even if the growth factor is no longer present. In the present model, external metabolic conditions such as glutamine levels are not represented. The model is designed to be extended to include metabolic effects on the cycle.

While the restriction point in the present model is in G1, recent research has questioned whether this holds for all cells [168], suggesting that that it may depend on cell type, and that cultured cells may have evolved with a shifted restriction point compared to wild-type cells [169]. The consensus remains that increasing Cyclin D-Cdk4/6 levels drive G1 progression into S, as in the present model, leaving the question of when in the cycle these levels are dependent on growth factors. The present model is consistent with the finding [170] that the restriction point coincides with the decline of APC^Cdh1^ before the G1-S transition.

In the present model, the inhibitory effect of actively replicating forks on new origin firing is applied directly to the rate of origin initiation as a damping term. While this damping was attributed to the ATR-Chk1 pathway, there are other mechanisms by which actively replicating forks could limit firing. Essential initiation factors such as Cdc45 are rate limiting [171], and are reintroduced only when forks terminate. Moreover, Chk1 has at least two mechanisms of damping initiation: it targets the essential initiation factor Cdc45 [172] and it degrades Cdc25A [173]. As it is unclear whether any of these mechanisms is the quantitatively dominant one, the damping term as in Eq. (7b) was used.

In the model, saturation of rate terms as functions of concentrations was only included where essential for proper model behavior. In actuality, all such terms must saturate, possibly beyond normal concentrations. Extension of the model to abnormal conditions may require inclusion of additional saturation effects.

The analysis of parameter sensitivity (Figure 4) implies that the model is relatively insensitive to changes in parameter values, in that cycling is maintained despite fivefold changes in parameters. Even so, these results indicate which parts of the model are more sensitive to parameter values. Cycling is inhibited if *k*_4_, *k*_11_, *k*_24_, *C*_8_ or *C*_10_ is doubled, or if *k*_9_, *k*_30_, *n*_3_, *C*_2_ or *C*_11_ is halved. Most of these parameters affect the levels and responses of SCF^βTrcp^, MPF, NUMA and securin-separase, the latter three of which control M phase. This suggests that mitosis may be more sensitive than the other phases to variations in levels of cell controllers or their reaction rates.

The chosen parameter values give a less abrupt fall of APCCdh1 at the G1-S transition than is suggested by the data for MCF-10A cells [174], which are non-cancerous but transformed mammalian cells [175]. If such behavior is found in untransformed cells, it could be modeled by increasing the Hill exponent n_2_ in Eq. (3) and also possibly changing the [SCF] term in Eq. (2) to have a Hill form. The model system shows sufficient flexibility to allow new sets of parameter values giving similar physiological behavior when such modifications are made.

Despite the different assumptions of Model Variants 1 and 2 regarding the licensing of origins, the two models give virtually the same results for all other variables, when equivalent parameter values are used. In the literature, some studies indicate that licensing occurs in G1 [176] while others state that it occurs in late M and G1 [177], or late M in tumor cells [95]. The model allows for these possibilities, and does not favor one or another.

This model uses a simplified representation of S phase, in which origins transit among four compartments with kinetics determined by cell cycle controllers and by the number of active origins, a surrogate for replication protein A, which senses single-stranded DNA. This model does not allow for local control of origin firing, for example in response to stalled forks [177]. Also, it does not explicitly represent the statistics of origin firing and the propagation speed of forks. It does not account for the fact that only a fraction of origins fire in cell cycle [178]. The present model provides a suitable framework for incorporating more detailed representations of S phase.

Mechanisms acting globally on origins, usually involving Cdk or geminin, are most often invoked as the primary ones preventing rereplication in mammalian cells. Here, it was noted that most of these mechanisms are unlikely to give an adequate level of prevention, with the cytoplasmic licensing factor being the lone exception. The key difference between this and other globally-acting mechanisms is that it gives a large temporal separation, relative to the cell cycle length, between the initial licensing step at nuclear membrane breakdown in M of the previous cycle, and origin initiation in S phase. The other plausible mechanisms act locally on origins. It is proposed here that global mechanisms will generally be inadequate unless they give such large temporal separation. The advantage of locally-acting mechanisms is that they obviate the need for temporal separation.

Concentration variables in the present model represent levels in the chromosomal region, and are assumed continuous in time throughout the cycle, including the processes of nuclear envelope breakdown and re-assembly. Because the substances considered have relatively high molecular weights, this is justified by the presence of the nuclear matrix which is highly structured, and observations that random thermal motions are not the primary determinants of transport inside cells [179].

In conclusion, a theoretical model for the mammalian cell cycle has been developed that satisfies the objectives listed in the Introduction. It differs significantly from previous models, being autonomous, continuous, cell-size independent, and largely based on experimentally observed mechanisms. It shows distinct behaviors corresponding to observed phases of the cell cycle, including a restriction point in response to external growth factors. The structure of the model makes it suitable for exploring the effects of the extracellular environment on cell cycle progression, including responses to metabolic conditions and to anti-cancer therapies that target progression of the cell cycle.

## Acknowledgements

This work was supported by NIH Grant T32 GM084905.

